# RBM10 variants and RBM5 involve in alternative splicing of RBM10v1 pre-mRNA: RBM10v1 includes its own exon 4, but RBM10v2 and RBM5 skip it

**DOI:** 10.1101/516930

**Authors:** Koji Nishio

**Author notes:** Address all correspondence to Koji Nishio Ph.D.

## Abstract

RNA binding motif (RBM) proteins, RBM10v1, RBM10v2 and RBM5 have quite similar molecular structures with a high degree of the conserved domains. Alternative splicing of RBM10 pre-mRNA produces the two mRNA variants, RBM10v1 (exon 4-included) and RBM10v2 (exon 4-skipped). RBM10v1 has a 77 amino acids-domain coded by its exon 4, but RBM10v2 lacks it. I explored the alternative splicing of the RBM10 pre-mRNA by the above three RBMs in COS-7, lung adenocarcinoma A549 and differentiated mouse cardiomyocytes H9c2 cells. Firstly, COS-7 and A549 cells express both RBM10v1 and RBM10v2 mRNA variants in contrast to H9c2 cells which express RBM10v2 variant alone. Transfection experiments of RBM10v1, RBM10v2 or RBM5 were performed to examine the alternative splicing of RBM10v1 pre-mRNA in COS-7, A549 and H9c2 cells. The result showed that RBM10v1 includes, by itself, its own exon 4 of the pre-mRNA in contrast to RBM10v2 and RBM5 which exclude the exon 4. The inclusion of the exon 4 seems to be repressed in differentiated H9c2 cells.

## 1. Introduction

Nuclear speckles is enriched in the pre-mRNA splicing regulators and locate in the interchromatin regions of mammalian cells (1). Splicing regulators are recruited to the active sites of transcription and the splicing machinery, spliceosomes. Stepwise assembly of various splicing regulators on the spliceosome is the most essential step. The initial step of splicing is binding of U1 snRNP with the pre-mRNA to form spliceosomal E complex (2). Binding of U2 snRNP to the branch point region leads to the formation of spliceosomal A complex. Preformed U4/U6/U5 tri-snRNP assembles with the precatalytic B complex and undergoes splicing of the target exon and intron. Proteomic studies of the complexes revealed the core protein composition and the interacting non-snRNPs which provided the molecular basis of inclusion or skipping of the alternative exons (3,4). The RNA binding proteins, HuR, PTB or TIA-1 antagonistically includes or skips the exon 6 of Fas receptor that encodes the transmembrane domain (5–7). In addition, RBM5 skips the exon 6 of Fas pre-mRNA in HeLa cells (8). Overexpressed RBM5 promotes distal alternative splice site pairing of the Fas pre-mRNA and thereby skips the alternative exon 6.

RBM5 modulates the expression of proapoptotic caspase 2L (9) and proapoptotic modulator Bax (10). In addition, RBM5 increases Stat5b and BMP5 mRNAs but decreases CDK2 and Amplified in Breast cancer-1, proto-oncogene pim-1 (a caspase antagonist) (11). In the breast carcinoma tissues, the expressions of two RBM10 variants and VEGF mRNAs significantly associate (12) and a similar association of RBM10v2, caspase 3 and proapoptotic Bax mRNAs is also reported (13). However, neither knockdown nor overproduction of RBM10 does not effect on alternative splicing of caspase 2 and 3 pre-mRNAs in HeLa cells (14).

Human RBM5 gene is frequently deleted in a variety of cancers (15) and RBM10 gene is also frequently mutated in lung adenocarcinomas (16). RBM10v1 (930 AA), RBM10v2 (852AA) and RBM5 (815AA) have significant sequence similarity as reviewed previously (17). So far, the functional relationship of the three RBMs has not been adequately clarified. The author has a hypothesis that the expression levels of the three RBMs possibly regulate alternative splicing of RBM10 pre-mRNA; accordingly, I explored a mechanism of alternative splicing of RBM10v1 pre-mRNA.

## 2 Materials and Methods

### 2.1 Cell lines and cell culture

H9c2 rat cardiomyoblasts, A549 and COS-7 cells were used. COS-7 cells were provided from JCRB (Japanese Cancer Research Resources Bank, Tokyo, Japan). COS-7 cells are SV40-large T antigen transformed immortalized cell line. COS-7 cells rapidly grew in the present study. COS-7 cells were used as a high expresser of RBM10 variants and RBM5. A549 human lung cancer cells were provided by Dr. Qiau (Chubu University, Kasugai, Aichi, Japan). A549 cells grew more slowly than COS-7 cells. A549 cells were used as a low expresser of RBM5 as previously described (10,18). H9c2 rat cardiomyoblasts cell line was provided by Dr. Tatsumi (Nagoya University). H9c2 cells have a potential to differentiate to cardiomyocytes (19). These cells were cultured in 6-well plates and passaged at a split ratio of 1:2 or 1:4. Each well contained 2 ml of Dolbeco’s modified essential medium (DMEM) supplemented with 10% FCS, 1 mM glutamine and 60 μg/ml kanamycine. The cells were grown until nearly full-confluent and the whole cell lysates were prepared (10 μg in SDS sample buffer) for Western blotting analysis of RBM10 and RBM5. The protein bands of RBM10V1, RBM10V2 or RBM5 were estimated by the image analysis software, Image-Pro plus (Media Cybernetics, MD. USA). Total RNA were prepared and used for RTPCR.

### 2.2 Antibodies and Western blot analysis

Rabbit polyclonal anti-RBM10 against the GST fused N-terminal 43 kDa region (#1-166) of rat RBM10v2 was provided from Dr. Inoue (Osaka City University) (20). The anti-RBM10 antibody specifically recognized the RBM10v1 (130 kDa) and RBM10v2 (114kDa) (20). The rabbit anti human RBM5 was provided by Dr. Oh (UCLA School of Medicine). The anti-RBM5 antibody was prepared against the COOH-terminal half of human RBM5 (amino acids 408–815) as described previously (21). The RBM10 and RBM5 proteins were successfully separated by SDS 8%-polyacrylamide gel electrophoresis based on their molecular sizes. The RBM proteins were identified with the primary rabbit anti-RBM10 or RBM5 antibody and the following secondary anti-rabbit IgG conjugated with Horse radish peroxidase (A6154; Sigma-Aldrich) or conjugated with alkaline phasphatase (T2180;Applied Biosystems).

### 2.3 Construction of expression vectors

An expression vector of wild type human RBM5-GPF was provided by Dr. Valcárcel (University Pompeu Fabra, Barcelona, Spain). An expression vector of human wild type RBM10v1 cDNA (930 amino acids) was provided from Kazusa DNA Research Center (Kisarazu, Chiba, Japan). Expression vectors of wild type rat RBM10v2 cDNA (852 amino acids), human RBM10v1-GFP and rat RBM10v2-GFP were provided by Dr. Inoue (Osaka City University, Osaka, Japan). RBM10 cDNA was amplified by polymerase chain reaction with high fidelity KOD DNA polymerase (Toyobo Inc., Japan). The coding sequences of the vectors or their protein expression were confirmed by cDNA sequencing, restriction digestion or western blotting of the protein with the specific antibodies

### 2.4 siRNA knockdown of RBM10

3’-Alexa Flour ^647^-labeled RBM10 siRNA pairs; sense, r(ACAGAUGGAUGGAAGCCAAdTdT and antisense, r(UUGGCUUCCAUCCAUGGUG)dTdA were obtained from QIAGEN (QIAGEN Japan, Tokyo). COS-7 cells grown at 80%-confluency in a 4-well culture slide were transfected with 50 pmoles of the RBM10 siRNA (at final concentration of 100 nM) by using 5 μl of Hyperfect siRNA transfection reagent. After 48 hours, the whole cell lysates were prepared and the protein levels of RBM10 and RBM5 were analyzed as described in the previous section. COS-7 cells subconfluently grown in 6-well plates were transfected with control siRNA or RBM10 siRNA, incubated for 48 hours and then total RNA and the cell lysates were prepared as described previously.

### 2.5 Reverse transcription and PCR analysis

Reverse transcription was performed from 1 μg of total RNA using Expand reverse transcriptase (Roche) and oligo dT or random hexamer oligonucleotides as a primer. PCR was performed with GeneAmp PCR System 9700 (Applied Biosystems) using FastStart Taq DNA polymerase (Roche) as described in the manufacturer’s instruction. The reaction mixture of PCR contained 1x PCR buffer, with or without 1x GC-RICH solution supplemented with 2 mM MgCl_2_, 0.2 mM of each dNTP, 0.5 μM of each forward and reverse primers and 2 Units of FastStart Taq DNA polymerase. cDNA was denatured at 95°C for 6min and then amplified for 35 cycles using the following parameters: 95°Cfor 30 sec, 55°C for 30 sec and 1min for 72°C, and at 72°C for 7 min (final extension). The endogenous mRNAs of human RBM10v1, RBM10v2, RBM5 and β-actin were amplified. To monitor the efficiency and reproducibility of PCR amplification, glycerolaldehyde 3-phosphate dehydrogenase (GAPDH), caspase 2L or β-actin cDNA was amplified as a control as described by Nishio et.al., (22).

For human RBM10v1 or RBM10v2, the forward primer RBM10A, 5’-AGGGCAAGCATGACTATGA-3’ and reverse primer RBM10B, 5’-GTGGAGAGCTGGATGAAGG-3’ were used as described by Martinez-Arribas et.al., (12) For human RBM5 (LUCA-15), the forward primer RBM5, LU15 (2) 5’-GACTACCGAGACTATGACAGT-3’ and reverse RBM5, LU15(3) 5’-AGAGGACAGCTGCACAAATGC-3’ 5’-GACTACCGAGACTATGACAGT-3’ were used as described by Rintala-Maki et.al (23). Forward primer RBM5A, 5’-GACTACCGAGACTATGACAGT-3’ and reverse primer RBM5B, 5’-AGAGGACAGCTGCACAAATGC-3’ were used as another primer pair. For human VEGF, the forward primer, 5’-AGCTACTGCCATCCAATCGC-3’ and reverse primer 5’-GGGCGAATCCAATTCC AAGAG-3’ were used as described by Wong et.al (24). For human caspase 2L (c-FLIP), the forward primer, 5’-GTTACCTGCACACCGAGTCACG-3’ and reverse primer 5’-GCGTGGTTCTTTCCATCTTGTTGGTCA-3’ were used according to Wang et.al (25). For human GAPDH, the forward primer, 5’-AAGGCTGAGAACGGGAAGCTTGTCATCAAT-3’ and reverse primer 5’-TTCCCGTCTAGCTCAGGGATGACCTTGCCC-3’ were used as described by Nishio et.al., (22). The amplified cDNA products were separated on agarose gels (1.8 % w/v) containing ethidium bromide (0.1 μg/ml). The cDNA were visualized by UV-exposure and the cDNA images were captured by Olympus **μ**-7000 digital camera.

## 3. Results

### 3.1. RBM10 siRNA inhibits expression of the endogenous RBM10 mRNAs, but not RBM5

COS-7 cells were transfected with the RBM10-siRNA and cultured for 48 hours. The whole cell lysates or total RNAs were subjected to western blotting analysis of the RBM proteins or RTPCR of the RBM mRNAs, respectively. The siRNA knockdown successfully down regulated the expression of RBM10v1 and RBM10v2, but not RBM5 as shown in Fig.1A. The results demonstrated that siRNA knockdown of RBM10 protein did not affect the expression of RBM5 mRNA. In addition, siRNA knockdown of the endogenous RBM5 under a similar condition was not successful due to a significantly high expression of RBM5 protein in COS-7 cells (data not shown).

Next I examined the effects of RBM10 siRNA and DOX on the alternative splicing of the Fas receptor mRNA as shown in Fig. 1B. The knockdown of RBM10 protein alone did not significantly alter the expression of two endogenous Fas receptor mRNA isoforms in COS-7 cells. Interestingly, the exposure to 1 or 2 μM DOX for 16 hours augmented the exon 6-skipped Fas 5-7 mRNA levels. In addition, the DOX-exposure decreased the RBM10 protein but not RBM5 (not shown). The DOX-exposed cells seemed to shift the equilibrium of the Fas isoform toward the exon 6-skiped Fas 5-7 mRNA due to the promotion of catalytic spliceosomes.

**Fig. 1.**
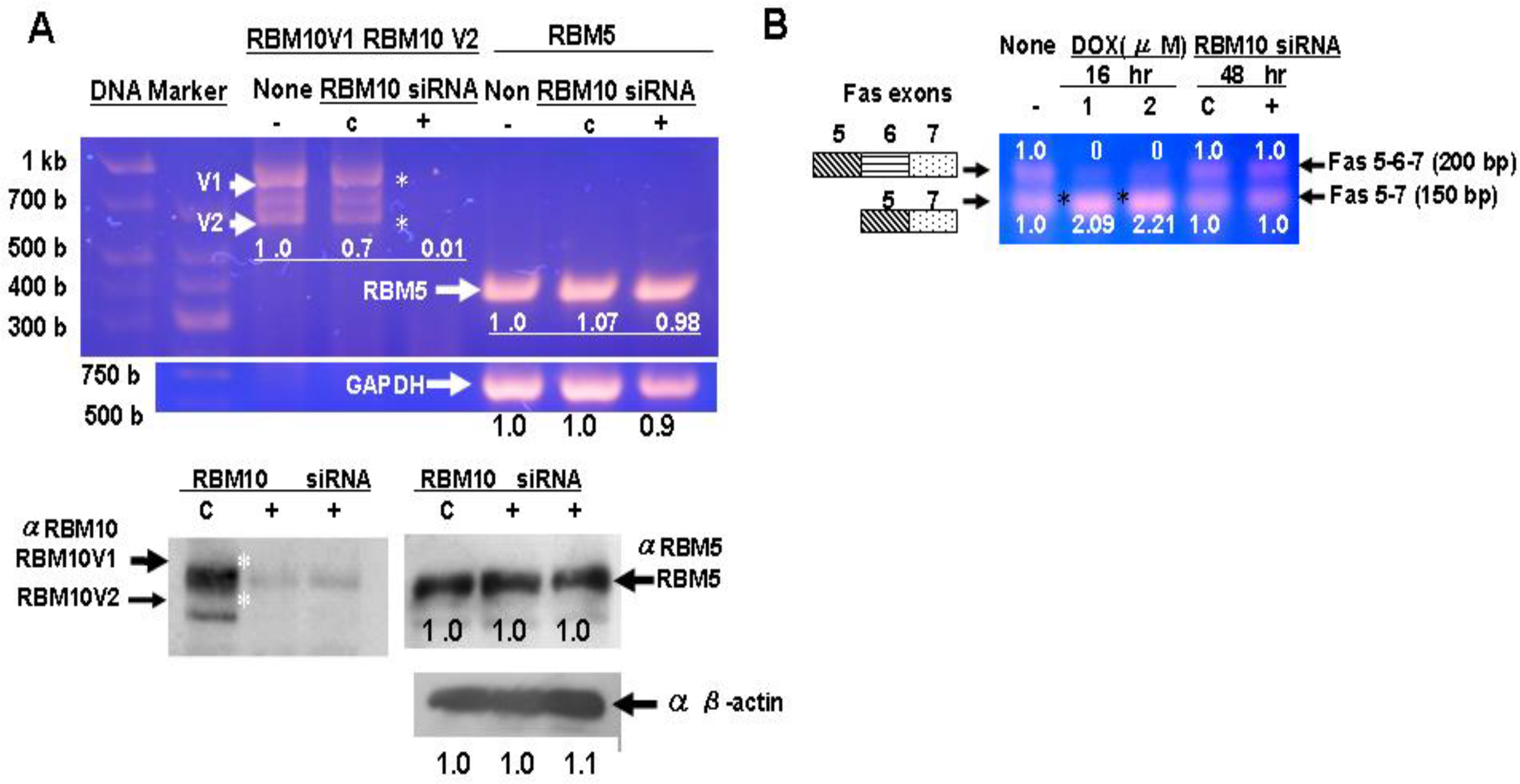
siRNA knockdown of RBM10 impaired the expression of RBM10 variants but not RBM5 in COS-7 cells. **A:** COS-7 cells grown in a 6-well plate were transfected with 100 nM RBM10 siRNA and then cultured for 48 hours or cultured without treatment (indicated as none). The total RNA of the non-treated cells (None), the cells treated with control siRNA (indicated as C), or those treated with RBM10 siRNA for 48 hours (+), was isolated and reverse-transcribed. The mRNA levels of RBM10v1, RBM10v2 or RBM5 were determined by RTPCR. The endogenous RBM10v1, RBM10v2, RBM5 or β-actin (as loading control) proteins were examined as described previously. The numbers below the amplified cDNA bands indicates the relative expression levels of the RBM10 or RBM5 mRNA. RBM10 siRNA inhibited the endogenous RBM10v1 and RBM10v2 expression, but not RBM5 and GAPDH mRNA (top panel). In a similar way, the siRNA significantly inhibited the expression of endogenous RBM10 protein, but not RBM5 (lower panel). **B:** COS-7 cells grown in a 6-well plate were transfected with RBM10 siRNA and cultured as described above, or cultured in the presence of 1 μM doxorubicin for 16 hours. The mRNA levels of Fas isoforms were analyzed by RTPCR as described above. The number above or below the each amplified cDNA band indicates the relative expression level of Fas (Fas 5-6-7: + exon 6) or Fas (Fas 5-7:- exon 6), respectively

### 3.2 RBM10v1 mRNA includes entire Exon 4: encodes particular 77 amino acids domain

RBM10v1 mRNA has a particular exon 4 which encodes 77-AA domain (amino acids: #68-144), whereas RBM10v2 lacks the exon 4. So far the function of the 77-AA domain has not been fully investigated. The sequence alignments of the exon 4 and exon 9 of human RBM5, RBM10v1 or RBM10v2 are shown in supplement S1. The 77-AA domain locates at amino terminal to the two RNA biding motifs. The region of #131-144 is included in the RNA recognition motif 1 as predicted by Interpro protein sequence analysis and classification (EMBL-EBI data base). The 77-AA domain consists of the unique amino acid sequences, an arginine-rich region: a 10 basic amino acids-cluster of Q<u>*RRRRRR*</u>H<u>*R*</u>H (#79-88), an aspartic acid-rich region: H<u>***R***</u>H***S***PTGPP***GF***PR<u>***D***</u>***G***<u>***D***</u>***Y***R<u>*D*</u>Q<u>***D***</u>***Y***<u>***R***</u> (#86-108) and a glutamic acid-rich region: a 13 acidic amino acids-cluster of <u>*E*</u>QG<u>*EEEEEEE*</u>D<u>*EEEEE*</u> (#110-125) and a part of RRM1 motif (IV***MLR***M***LP***QAA***TE***D***D***). The conserved amino acids are highlighted in italic bold. The basic and acidic residues are shown in underlined italic. The above-mentioned arginine and glutamic acid repeats, respectively exhibit strong positive and negative charge distribution which may suggest a potential biological role. Thereby, I termed the unique domain as ***RDE-rich*** *domain* of RBM10v1.

RBM5 has its own exon 4 which encodes 52 amino acids (#62-113) consisting of an aspartic acid-rich region (R***R***N***S***<u>*D*</u>RSE<u>*D*</u>***GY***HS<u>***D***</u>*G*<u>***D***</u>*Y*GEH<u>***D***</u>***YR***) and a part of the RRM1 motif (TI***MLR***G***LP***ITI***TE***S***D***). The exon 4 of RBM5 lacks most of the arginine and glutamic acid repeats. Consequently the common structural features of RBM10v2 and RBM5 are lacks of the positive and negative charge clusters: Q*RRRRRR*H*R*H and <u>*E*</u>QG<u>*EEEEEEE*</u>D<u>*EEEEE.*</u> RBM5 has its own exon 9 which encodes only 22 amino acids, but RBM10v1 and RBM10v2 have exon 9 which encodes 59 amino acids. The 59-AA domain locates at amino terminal to the highly conserved RRM2 that encoded by exon 10 and exon 11 (not indicated). The function of the 59-AA domain is unknown at this moment. After all, the ***RDE-rich*** domain seems to have a particular role in the formation of pre-catalytic spliceosomal complex with the target pre-mRNA.

Alternative splicing factors which involve in inclusion or skipping of the exon 4 of RBM10 pre-mRNA have not been thoroughly investigated. In the present study, I explored a role of the ***RDE*-rich** domain of RBM10v1 with regard to the inclusion of its own exon 4.

### 3.3 RBM10v1 includes the alternative exon 4 of RBM10v1 pre-mRNA, but RBM10v2 and RBM5 skip it in COS-7 cells

Firstly, COS-7 cells grown in wells of a 6-well plate were transfected with4 *μ*g of the expression vector, pRBM10v1-EGFP, pRBM10v2-EGFP or pEGFPN1 (vector alone), and then cultured for the following 48 hours. The sufficient protein expression of RBM10v1-GFP or RBM10v2-GFP could be confirmed by observing the green fluorescent nuclear speckles. The expression levels of RBM10v1, RBM10v2 and RBM5 mRNAs of the transfected cells were analyzed by RTPCR. The representative data were shown in Fig. 2A and the bar graph was displayed in supplement Fig S2A.

**Fig. 2.**
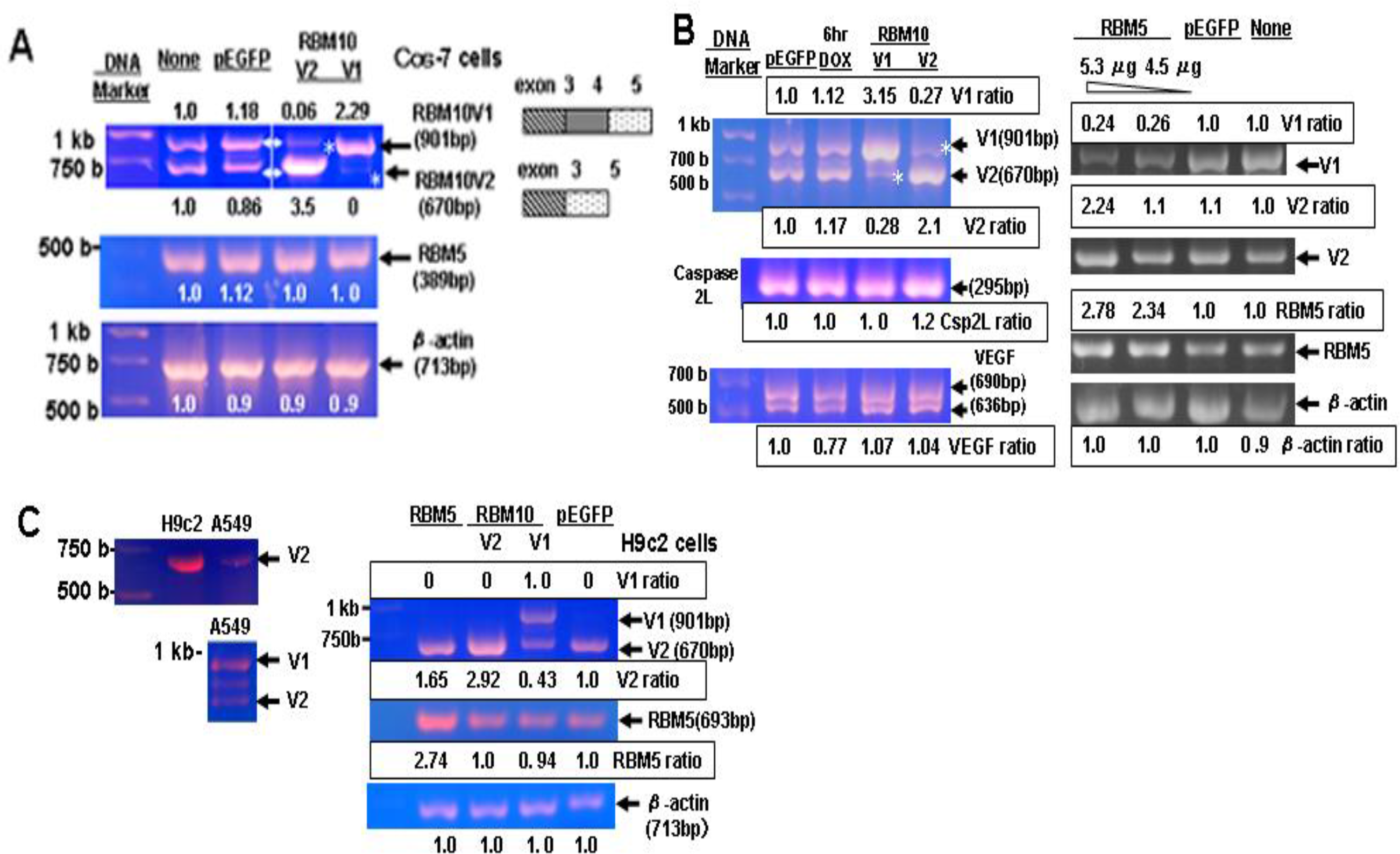
Inclusion or skipping of exon 4 of RBM10v1 is caused by expression of RBM10v1-GFP or RBM10v2-GFP in COS-7cells. (**A**) COS-7 cells subconfluently grown in 6-well plates were cultured without treatment (none) or transfected with 4 μg of pEGFPN1, pRBM10v1-GFP or pRBM10v2-GFP vector and cultured for the following 48 hours. The none-treated cells (None) or pEGFP transfected cells (pEGFP) were used as control. The mRNA levels of the endogenous RBM10v1, RBM10v2, RBM5 or β-actin were analyzed by RTPCR and calculated by Image-Pro Plus software. The numbers above or below the each amplified cDNA bands indicate the relative expression levels of RBM10v1, RBM10v2, RBM5 or β-actin mRNA. (**B**) Exon 4-inclusion or skipping of RBM10v1 pre-mRNA promoted by RBM10v1-GFP or RBM10v2-GFP in A549 cells. A549 cells subconfluently grown in 6-well plates were treated with 1 μM doxorubicin for 6 hours, or transfected with pEGFPN1 (as a control), pRBM10v1-GFPor pRBM10v2-GFP and cultured for 48 hours (left panel). A549 cells were transfected with 4*μ*g of pEGFPN1 or 4.5 or 5.4 `μ`g of pRBM5-GFP and cultured for 48 hours (right panel). The mRNA levels of the endogenous RBM10v1, RBM10v2, RBM5, VEGF, caspase 2L or β-actin were analyzed by RTPCR. None-treated cells (not included) gave a quite similar result as the pEGFP transfected cells. The endogenous caspase 2L mRNA levels (also as loading control) were quite similar revels. A pair of the RBM5A and RBM5B primers was used for amplification of the RBM5 mRNA (693 bps). The numbers above or below the each amplified cDNA bands indicate the relative expression levels of RBM10v1, RBM10v2 or RBM5 mRNA. (**C**)H9c2 cells express mostly RBM10v2 (left panel). H9C2 cells were transfected with 4 `μ`g of pEGFP (as a control), pRBM10v1-GFP, pRBM10v2-GFP or pRBM5-GFP as described above. The expression of RBM10v1, RBM10v2, RBM5 or β-actin mRNA levels were examined as described in **B**. The pEGFP-transfected H9c2 cells expressed mostly endogenous RBM10v2 mRNA alone (left panel). The endogenous β-actin mRNA levels (as loading control) were quite similar revels. The pRBM5-GFP transfected cells produced 2.74- fold more RBM5 mRNA than the pEGFP, pRBM10v1-GFP or pRBM10v2-GFP transfected cells. The exogenous expression of RBM10v1-GFP decreased the endogenous RBM10v2 mRNA level to about 43% of the control level.

The non-treated and pEGFPN1-transfected Cos7 cells expressed the quite similar levels of the endogenous RBM10v1 mRNA (the relative levels of 1.0 and 1.18) and those of the endogenous RBM10v2 mRNA (1.0 and 0.86), respectively. When the exogenous RBM10v2-GFP was expressed, the endogenous RBM10v1 mRNA was significantly impaired to 0.06, but the RBM10v2 mRNA increased more than threefold (the relative level of 3.5) as shown in Fig. 2A, top panel and supplement Fig. S2A. More than 94 % of the endogenous RBM10v1 mRNA seemed to be exon 4-skipped by the exogenously expressed RBM10v2-GFP. In vice versa, the exogenously expressed RBM10v1-GFP impaired the endogenous RBM10v2 mRNA to the undetectable level, but RBM10v1 mRNA increased more than twofold (the relative level of 2.29). These results suggest that the exogenous RBM10v1 includes the exon 4 of the endogenous RBM10 pre-mRNA, whereas the exogenous RBM10v2 skips it. Hence, the inclusion of the exon 4 promoted by RBM10v1 depends on the ***RDE-rich*** domain and the skipping of the exon 4 promoted by RBM10v2 regards as antagonistic. In addition, I made the N-terminal 481 residues-truncated RBM10v2 construct (#482-852) that lacks the entire RRMs. The expression of the truncated RBM10v2 did not affect the endogenous RBM10v1 and RBM10v2 mRNA levels (data not shown).

### 3.4 Alternative splicing of RBM10v1 pre-mRNA in A549 lung carcinoma cells

Next, I examined the alternative splicing of RBM10 pre-mRNAs in A549 lung cancer cells (a low expresser of RBM10 and RBM5) as shown in Fig. 2B, left panel and supplement Fig S2-B. The expression of RBM10v1-GFP increased the RBM10v1 mRNA more than threefold but decreased the endogenous RBM10v2 mRNA to 28 % of the control level. The expression of RBM10v2-GFP increased the RBM10v2 mRNA more than twofold but decreased the endogenous RBM10v1 mRNA to 27 % of the control level. Again the results reconfirmed that the inclusion or skipping of the exon 4 is promoted by the expression of RBM10v1or RBM10v2, respectively.

The EGFP-expressed A549 cells expressed the proapoptotic caspase 2L mRNA, but a little caspase 2S mRNA (precursor of caspase 2L, not shown). Thereby the caspase 2L mRNA was not additively increased by the DOX treatment. The caspase 2L and the VEGF mRNAs were not affected by RBM10 variant expression, and thereby they were not alternatively spliced by RBM10 in A549 cells. Therefore, the caspase 2L mRNA was used as an internal control for the mRNA levels. It is known that RBM10 did not involve in caspase 2L pre-mRNA alternative splicing in HeLa cells also (14). The short exposure of DOX to A549 cells for 6 hours did not alter the equilibrium of RBM10 mRNA variants. The tumor specific VEGF mRNAs were also not significantly affected by DOX.

### 3.5 RBM5 skips exon 4 of RBM10v1 pre-mRNA in A549 cells

Tumor suppressor RBM5 was examined for the involvement in alternative splicing of RBM10v1 pre-mRNA. A549 cells were transfected with pRBM5-GFP and the changes of RBM10 mRNA variants were analyzed. The non-treated cells expressed the two RBM10 mRNA variants at similar levels. In the pRBM5-GFP transfected cells, the total RBM5 mRNA and the endogenous RBM10v2 mRNA increased threefold and more than twofold, respectively, in contrast the endogenous RBM10v1 mRNA decreased to less than 24 to 26 % of the control as shown in Fig. 2B, right panel and supplement Fig. S2-C. The endogenous RBM10v1 mRNA expression inversely correlated with the higher expression levels of RBM5 (see the ratio of v1/RBM 5 in Fig.S2-E) and the higher expression of RBM5 increased the endogenous RBM10v2 mRNA levels (see the ratio of v1/v2). The result indicates that the exogenous expression of RBM5 specifically decreased the endogenous RBM10v1 mRNA and thereby increased the endogenous RBM10v2 mRNA. A549 cells expressed the abundant caspase 2L mRNA but a little caspase 2S mRNA, a precursor of caspase 2L. In fact, the detection of a minor increase of the caspase 2L mRNA was difficult in the RBM5-GFP expressed A549 cells.

### 3.6 H9c2 cells express RBM10v2 alone but not RBM10v1

H9c2 cardiomyoblasts predominantly expressed RBM10v2 mRNA as shown in Fig. 2C, left panel. Next, H9c2 cells were transfected with pEGFP, pRBM10v1-GFP, pRBM10v2-GFP or pRBM5-GFP and the following expression levels of RBM10v1, RBM10v2 or RBM5 mRNA were compared by RTPCR as described previously. The pEGFP-transfected H9c2 cells predominantly expressed the endogenous RBM10v2 mRNA as shown in Fig 2C, right panel and supplement Fig S2-D. The exogenous expression of RBM5-GFP and RBM10v2-GFP significantly increased the RBM10v2 mRNA levels nearly 1.7-fold and threefold, respectively, due to the additional production of the endogenous or exogenous RBM10v2 mRNA. When the exogenous RBM10v1-GFP was expressed in H9c2 cells, the endogenous RBM10v2 mRNA decreased to 43% of the control level. In addition, the expression of RBM10v1-GFP and RBM10v2-GFP did not affect the endogenous RBM5 mRNA levels (supplement Fig.S2-D). Finally the present investigation demonstrated that the alternative splicing of RBM10 pre-mRNA is regulated by RBM10v1, RBM10v2 and RBM5 in COS-7 cells, A549, and H9c2 rat cardiomyoblasts.

## Discussion

### siRNA knockdown of RBM10 does not alter the expression of RBM5 mRNA and Fas receptor mRNA variants in COS-7 cells

A preliminary analysis indicated that COS-7 cells expressed a several-fold higher amount of RBMs than HeLa and A549 cells. In the present study, I performed siRNA-knockdown of RBM10 in COS-7 cells which represent a highest expresser of RBMs. The successful RNA interference of RBM10 did not affect the expression levels of RBM5 mRNA and protein. In addition, the siRNA-knockdown of RBM10 did not affect the expression of the Fas 5-6-7 (+exon 6) and Fas 5-7 (-exon 6) mRNAs. In COS-7 cells, the exogenous expression of RBM5 significantly promoted exon 6 skipping of Fas-pre-mRNA as shown in supplement Fig.S1, in contrast those of the exogenous RBM10v1 and RBM10v2 which did not alter the two Fas isoform levels. Interestingly, the exposure of 1 μM or 2 μM DOX for 16 hours significantly decreased the Fas receptor mRNA (Fas 5-6-7), and thereby the Fas 5-7 mRNA increased about twofold. Previously I found that the DOX-treated cells significantly decreased their RBM10 protein but not RBM5. Therefore the activated RBM5 of the DOX-treated cells seemed to promote the expression of Fas 5-7 mRNA isoform. Taken together, these results suggest that the major variant RBM10v1 and RBM10v2 do not promote the alternative splicing of Fas pre-mRNA, because the actively growing COS-7 cells express a higher amount of RBM5 protein at stationary phase.

### Antagonistic Function of RBM10v1, RBM10v2 and RBM5 with regard to alternative splicing of exon 4 of RBM10v1 pre-mRNA

COS-7 and A549 cells expressed RBM10v1 and RBM10v2. However H9c2 cardiomyoblasts expressed RBM10v2 alone. This finding suggests a differential regulatory mechanism to balance or shift the equilibrium between RBM10v1 and RBM10v2 in the highly proliferative cells or differentiated cells. Hence, I explored the switching between the two RBM10 variants in COS-7 cells (high expresser of RBMs), A549 cells (moderate expresser of RBMs) and H9c2 cells (low expresser of RBMs). The cells were transiently transfected with pRBM10v1-GFP or pRBM10v2-GFP construct. The exogenous expression of RBM10v1-GFP predominantly increased the RBM10v1 mRNA but significantly impaired the endogenous RBM10v2 mRNA. The expression of RBM10v2-GFP also increased the RBM10v2 mRNA but impaired the endogenous RBM10v1 mRNA. The result indicates that RBM10v1 specifically recognizes its own exon 4 of RBM10v1 pre-mRNA and facilitates its inclusion, whereas RBM10v2 skips it.

Some other splicing regulator can skip the alternative exon of its own pre-mRNAs. One example is polypyrimidine tract binding protein (PTB) which has an autoregulatory activity. Overexpression of PTB activates the skipping of exon 11 from its own pre-mRNA, yielding a premature termination codon containing isoform that are targeted to nonsense-mediate decay (26). In addition, RBM4 skips the exon 11 of PTB pre-mRNA to down-regulate PTB, as myoblasts commit to terminal differentiation (27).

### Alternative splicing mechanism of RBM10 pre-mRNA by RBM10v1, RBM10v2 and RBM5

RBM10v1 and RBM5 mRNA has its own exon 4 which encodes 77 and 52 amino acids, respectively, whereas RBM10v2 lacks the entire exon 4 as shown in supplement S1. Exon 4 of RBM5 encodes an aspartic acid-rich region and a part of RRM1, but lacks the arginine-rich (DSYEASPGSETQRRRRRR) and glutamic acid rich (EEEDEEEEEKA) regions. The present investigation demonstrates that RBM10v2 and RBM5 have a distinct potential for involving the skipping of the alternative exon 4 of RBM10v1 pre-mRNA. The above unique molecular structure and the highly conserved RNA recognition motifs (RRM) appear to associate with a distinct binding specificity of the target pre-mRNAs as mentioned below. RBM10v1, RBM10v2 and RBM5 have RRM1 and RRM2 as shown in supplement S2. The amino acid sequence identities of the RRMs between four RBM proteins: RBM10v1, RBM10v2, RBM5 and RBM6 were calculated by the alignment program (Vector NTI Suite 7.0 AlignX) as shown in supplement S2. To estimate the positive identities of the RRMs, the amino acid substitutions among isoleucine, valine, methionine or leucine, those among threonine, serine or alanine or those among aspartic and glutamic acid were calculated as the conserved amino acids. RRM1 and RRM2 of RBM10v1 share the identities of 98.4 % and 98.8% with those of human RBM10v2. The two RRMs of RBM10v1 share the identities of 59.5 % and 69.5% and in addition higher positive identities of 68.4% and 87.8% with those of human RBM5. The two RRMs of human RBM10v1 share the lower identities of 29.5% and 32.9% and lower positive identities of 44.9% and 56.1% with those of RBM6. After all RBM10v1, RBMv2 and RBM5 share the higher identities of the RRMs.

Interestingly V354E-mutation in the RRM2 disrupts the function of RBM10 and does not inhibit proliferation of A549 lung adenocarcinoma, because the mutated RBM10 does not promote skipping of exon 9 of the gene NUMB (28,29). Recently, the solution structure of the RBM5 RRM2 core domain was determined and its flexible RNA binding properties were revealed (30). The RBM5 RRM2 can preferentially bind both CU and GA rich sequences (5’-CUCUUC-3’ or 5’-GAGAAG-3’). Taken together it is likely that the flexible RNA binding properties of RBM5 and RBM10 and in addition a unique ***RDE-rich*** domain of RBM10v1 seem to involve with alternative splicing of RBM10 pre-mRNA.

Then, I explored the expression of the RBM10v2 mRNA (exon 4-skipped RBM10) and its promotion by the exogenously expressed RBM5 in A549 cells (moderate expresser of RBM5) and H9c2 cells (expresser of RBM10v2 and RBM5). Interestingly the RBM5 expression predominantly decreased the endogenous RBM10v1 mRNA in A549 cells but increased the endogenous RBM10v2 mRNA in A549 cells or H9c2 cells. The ratios of v1/v2 and v1/RBM5 were decreased in the RBM5-GFP-transdected, but not in the none-treated and pEGFP-transfected cells as shown in supplement Fig S2E. The results suggest that among the high or low expressers of RBMs, the activities of RBM10v1, RBM10v2 or RBM5 may set a unique balance of the RBM10v1 and RBM10v2 proteins.

RBM5 plays a key role as an alternative splicing regulator of the proapoptotic factors as described previously (9,10). A molecular mechanism of alternative splicing of Fas receptor pre-mRNA is based on the pre-spliceosomal assembly of RBM5 (8). RBM5 directly interacts with exon 6 of the Fas pre-mRNA on the pre-spliceosome and does not affect the association of U1 and U2 snRNPs to the adjacent splice sites. Finally RBM5 inhibits the incorporation of the U4/U6-U5 tri-snRNP complex on the introns flanking the Fas-exon 6, and promotes the pairing of U1 and U2 at the distal splice sites, contributing to skipping the exon 6.

Sixty amino acids-OCRE domain of RBM10v1 (#561-621) and RBM5 (#451-511) has the OCRE sequence motif that contains a unique tyrosine-rich sequence (31–33). The OCRE domain is highly conserved among human RBM10v1, RBM10v2 and RBM5 (with the positive identities of 100 %). Mutation of the conserved aromatic residues of the OCRE-motif impaired the skipping of Fas exon 6 (31). Thereby the OCRE-motif is required for alternative splicing after targeting the RBM10v1 pre-mRNA.

The present investigation indicates that exon 4 skipping of RBM10v1 is similar in mechanism of exon 6 skipping of Fas by RBM5. It is intriguing to speculate that RBM10v2 and RBM5 interact with the RBM10v1 exon 4 and the flanking introns without affecting association of U1 and U2 snRNP to the splice sites. RBM10v2 and RBM5 can promote the pairing of U1 and U2 at the distal splice sites (Fig. 3B). Notably RBM10v1 involves in inclusion of its own exon 4. Thereby, RBM10v1 must interact with the exon 4 and stimulates the incorporation of the U4/U6-U5 tri-snRNP complex on the introns flanking exon 4. RBM10v1 promotes the pairing of U1 and U2 at the proximal splice sites, contributing to the exon 4-inclusion (Fig. 3A). Therefore, the ***RDE-rich*** domain of RBM10v1, its arginine-rich or glutamic acid-rich regions are indispensable for the inclusion process. The lack of both regions of RBM5 and RBM10v2 suggests a particular role of the ***RDE-rich*** domain of RBM10v1. The present results revealed the antagonistic function among the RBM10v1 and RBM10v2 or RBM5, thereby alternatively generating the two RBM10 variant mRNAs. I did not clarify direct binding of the RBM10 pre-mRNA with RBM10v1, RBM10v2 or RBM5 protein. Their direct interaction with the RBM10-exon 4 and flanking introns should be demonstrated in the future study.

**Fig. 3.**
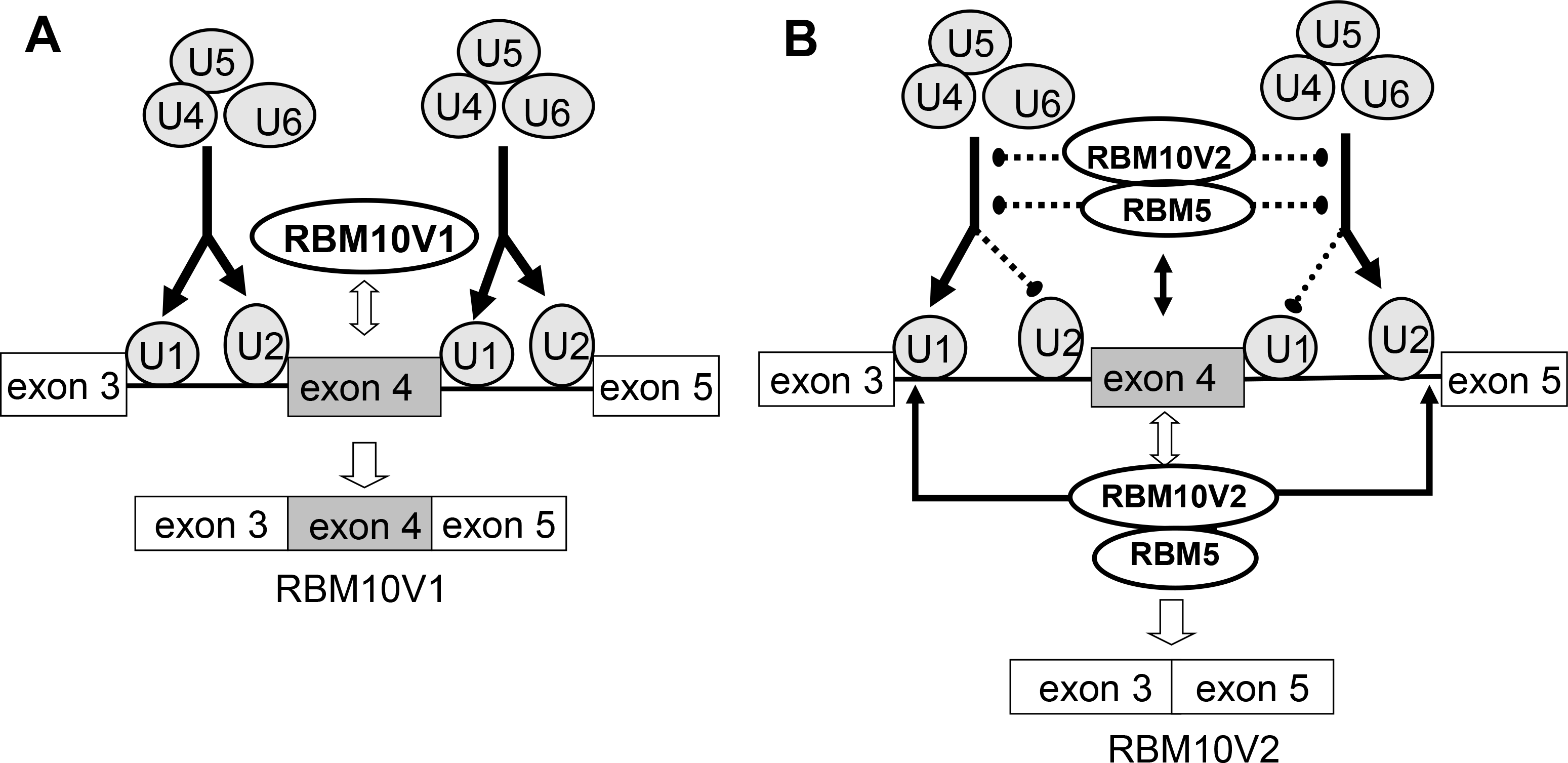
An antagonistic model for RBM10v1 pre-mRNA alternative splicing regulated by RBM10v1, RBM10v2 or RBM5. The constitutive exons are shown as white boxes, and the alternative exon 4 is shown as a gray box. The intron 3 or intron 4 is shown by a thin line. (**A**) RBM10v1 includes its own alternative exon 4 by binding of the U4/U5/U6 tri-snRNP on the pre-spliceosomal complex in the flanking introns of the exon 4. RBM10v1 promotes the pairing of U1 and U2 snRNP as the proximal splice sites, contributing to the inclusion of the exon 4. (**B)** RBM10v2 or RBM5 skips the exon 4 of RBM10v1 pre-mRNA by blocking of the U4/U5/U6 tri-snRNP at the proximal splice sites on the pre-spliceosomal complex. The dotted line shows blocking of the proximal site paring. Thereby, RBM10v2 or RBM5 promotes the pairing of U1 and U2 as the distal splice sites to skip the alternative exon 4.

### Correlation of RBM10 variant and other RBMs in human cancer cells or tumor tissues of the advanced grade

Differential RTPCR analysis of the X-chromosome RBM gene expressions was performed in human breast carcinoma tissues, which indicated that the expressions of RBM10v2 associate with high proliferation of the tumors and those of RBM10v1 and RBM10v2 are interdependent (12). Notably, in the breast carcinoma tissues, a significant inverse correlation of the mRNA levels between the breast cancer metastasis suppressor-1 (BRMS1) and RBM10v1 was reported (34).

Correlated expressions between RBM5 and RBM10v1 or those between RBM6 and RBM10v2 are highly significant in the breast tumor tissues (35). The positive correlation between RBM6 and RBM10v2 mRNA increases in relation to the advanced tumor grade, increased tumor size and loss of progesterone receptor. The two RBM mRNA expressions are highly significant in the Grade 3 ductal carcinoma. RBM5, RBM10v1 and HER2 protein expression levels are elevated in the tumor tissues. The consistent strong positive correlations of the RBMs may suggest their functional interrelation or antagonism. RBM5 modulates the expression of proapoptotic factor Bax, enhances caspase-3 activity and causes apoptosis in A549 lung cancer cells (10). In addition, RBM10v2, Bax and caspase 3 mRNAs are coexpressed in the breast tumor tissues (13). The statistically significant correlation of the above three proteins suggest that the both of RBM5 and RBM10v2 may involve in the splicing of the Bax pre-mRNA.

So far, molecular mechanism of the interdependency or correlation among the RBMs has not been clarified. In the present study, I found the inverse interdependency of RBM10 variants and the positive correlation of RBM5 and RBM10v2 mRNA. It is intriguing to speculate that the inverse interdependency of the RBM10 variants may be associated with accompanying alternative splicing of RBM10 pre-mRNA by RBM10 variant itself and RBM5. The primarily generated RBM10v1 will increase its own RBM10v1 mRNA. In contrast, RBM5 has a potential for generating the RBM10v2 variant. The secondary generated RBM10v2 will increase its own mRNA variant and the translation product, RBM10v2 that facilitates its own mRNA production further. Therefore, significantly high expression of RBM10v1 may reflect its antagonistic function to RBM10v2 and RBM5. Finally, RBM10v1 variant may be dominantly expressed in a variety of cancers that deleted RBM5 gene.

A previous investigation revealed a nonsense mutation (at 1235G>A) or a flame shift mutation (at c1893_1894insA) of human RBM10 gene (36). The mutated RBM10 gene seems to produce a truncated RBM10v2 protein (411 or 632 amino acids) or truncated RBM10v1 (488 or 709 amino acids) that remains the RRM motifs. In the affected families, the nonsense or null mutations in human RBM10 gene, causes TARP syndrome, talipes equinovarus, atrial septal defect, Robin sequence and persistent left superior vena cava. Whole exon sequencing /genome sequencing of 183 lung adenocarcinoma/normal pairs revealed that frequent somatic mutations in an epigenetic regulator/ tumor suppressor and splicing regulators (U2AF1, RBM10, ARIDA1, SETD2 and BRD3). Two splice-site or 5 truncating mutations are found in the RBM10 gene of the lung adenocarcinomas (16). The alterations of these genes are speculated as a novel hallmark of epigenetic and RNA deregulation. Finally the two RBM10 variants and RBM5 will involve with alternative splicing of the possible target pre-mRNAs, the products of which seem to participate in the embryonic development and cancer prevention of the normal tissues.

## Acknowledgements

This work was supported in Grant in Aid for Science Research (C) by the Japan Society for the Promotion of Science (to K.N.; 17590159).

## Author Contributions

K.N conceived, designed and performed the experiments, and wrote the manuscript.

**Supplement S 1** Alignments of the sequences coded by exon 4, exon 9 and exon 10 of human RBM5, RBM10v1 or RBM10v2. The identical residues in the all sequences are red in the highlighted yellow columns. The consensus residue derived from a block of similar residues at a given position among the two RBM is blue in the highlighted light blue columns. The consensus residue derived from the occurrence of greater than 50% of a single residue at a given position is black in the highlighted green columns. The unique residues of RBM10v1 exon 4 are black in the highlighted yellow columns. The non-similar residues are black. The dotted lines show the lack of the corresponding residues. A double headed arrow above the alignment shows a part of RRM1 or RRM2 coded by exon 4 or exon 10, respectively.

**Supplement S2** Alignments of RRM1 and RRM2 motifs of RBM10v1, RBM10v2, RBM5 or RBM6. The consensus residues derived from a completely conserved residue at a given position among all the RRM motifs are red in the highlighted yellow columns. The consensus residue derived from a block of similar residues at a given position among the RRM motifs is blue in the highlighted light blue columns. The consensus residue derived from the occurrence of greater than 50% of a single residue at a given position among the motifs is black in the highlighted green columns. The residue weakly similar to consensus residue at given position is green. The non-similar residues are black. The identity or positive identity of each RRM motif is indicated.

**Supplement Fig. 1.** Skipping of Fas exon 6 by treatment with DOX or expression of RBM5-GFP in COS-7 cells. COS-7 cells subconfluently grown in 6-well plates were cultured without treatment (as a control: None) or transfected with 4 μg of pEGFPN1, pRBM10v1-GFP, pRBM10v2-GFP or pRBM5-GFP vector and cultured for the following 48 hours. COS-7 cells were cultured with 1 μM DOX for 6 hours. The mRNA levels of the endogenous Fas 5-6-7 and Fas5-7 were analyzed by RTPCR as previously described in the legend of figure 1. Asterisk (*) denotes the significant production of Fas5-7 isoform.

**Supplement Fig.2**. Analysis of the mRNA levels of RBM101, RBM10v2 and RBM5 in COS-7, A549 and H9c2 cells. The cells grown in 6-well plates were cultured without treatment (as control: None) or transfected with the expression vectors as described in figure 2. The mRNA levels of the endogenous RBM10v1, RBM10v2 and RBM5 and those of the exogenously expressed RBM were analyzed and displayed as bar graphs. COS-7 cells (A), A549 cells (B and C), and H9c2 cells (D). Caspase and β-actin mRNAs were employed as loading control. A549 cells were transfected with RBM5-GFP and the endogenous RBM10v1 and RBM10v2 were investigated (C) and the ratio of RBM10v1/RBM10v2 or RBM10v1/RBM5 were displayed (E).

