## supplement S1 for "RBM10 variants and RBM5 involve in alternative splicing of RBM10v1 pre-mRNA: RBM10v1 includes its own exon 4, but RBM10v2 and RBM5 skip it"

```
                                                                                                  part of RRM1
                                                                                              <---------------->
                  68         79       88                   108             125  131         144
RBM5 exon 4      RERERR--------------------NSDRSEDGYHSDGDYGEHDYRHDISDERE-----------SKTIMLRGLPITITESD
RBM10v1 exon 4         DSYEASPGSETQRRRRRRHRHSPTGPPGFPRDGDYRDQDYRTEQGEEEEEEEDEEEEEKASNIVMLRMLPQAATEDD
RBM10v2                -----------------------------------------------------------------------------

RBM5 exon 9      DSEQEVPPGTT-------------------------------------ESVQSVDYYCD
RBM10v1 exon 9   EAEQKLPLGTRLDQQTLPLGGRELSQGLLPLPQPYQAQGVLASQALSQGSEPSSENAND
RBm10v2 exon 9   EAEQKLPLGTRLDQQTLPLGGRELSQGLLPLPQPYQAQGVLASQALSQGSEPSSENAND

                                  part of RRM2
                  <-------------------------------------------------->
RBM5 exon 10     TIILRNIAPHTVVDSIMTALSPYASLAVNNIRLIKDKQTQQNRGFAFVQLSSAM
RBM10v1 exon 10  TIILRNLNPHSTMDSILGALAPYAVLSSSNVRVIKDKQTQLNRGFAFIQLSTIV
RBM10v2 exon 10  TIILRNLNPHSTMDSILGALAPYAVLSSSNVRVIKDKQTQLNRGFAFIQLSTI-
```
