## Supplementary figures and images for "RBM10 variants and RBM5 involve in alternative splicing of RBM10v1 pre-mRNA: RBM10v1 includes its own exon 4, but RBM10v2 and RBM5 skip it"

### supplement Fig S1

Supplement Fig. S1

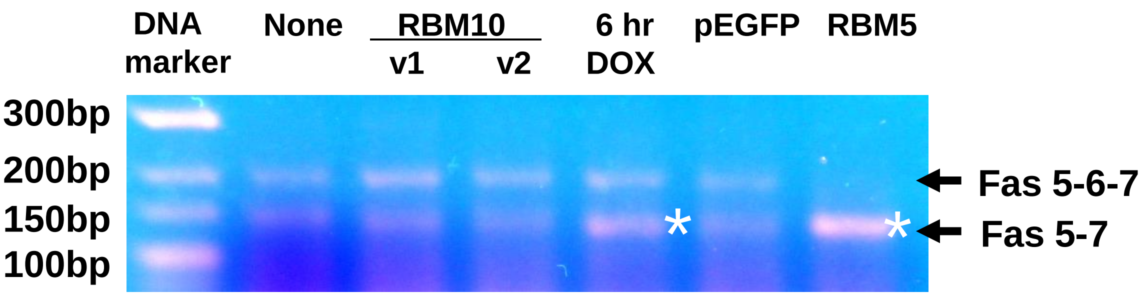


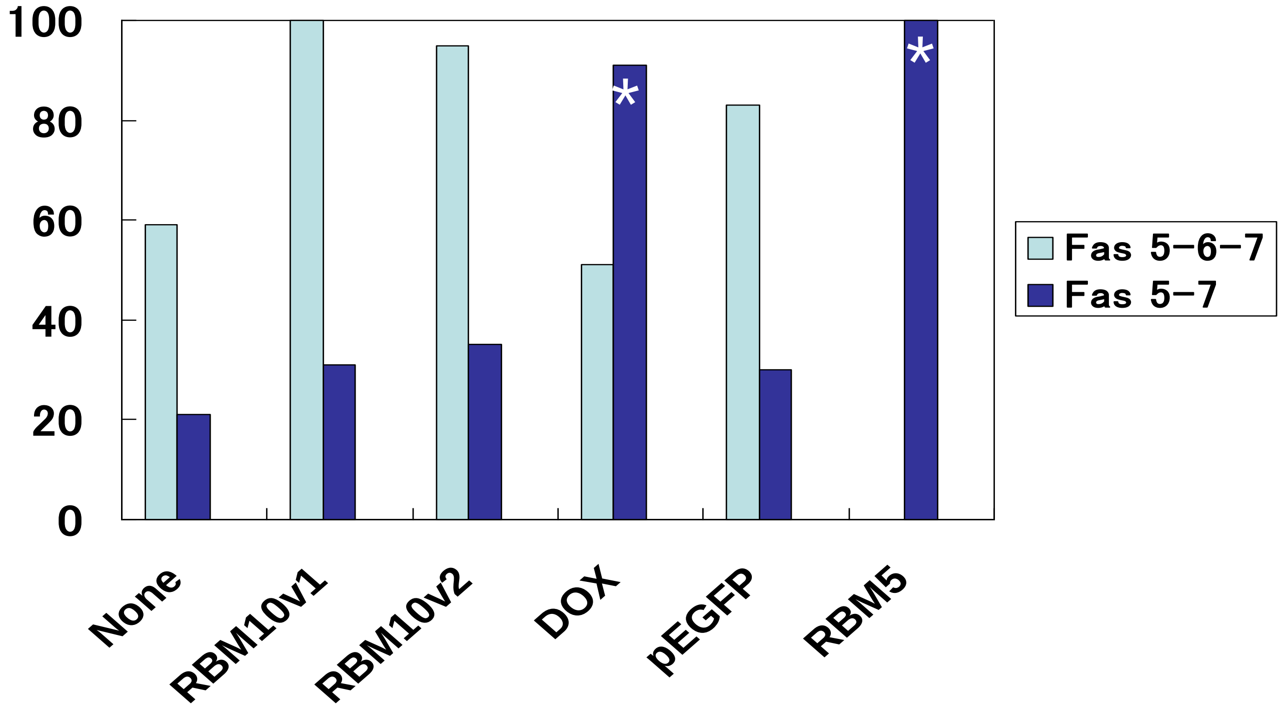

### supplement Fig S2

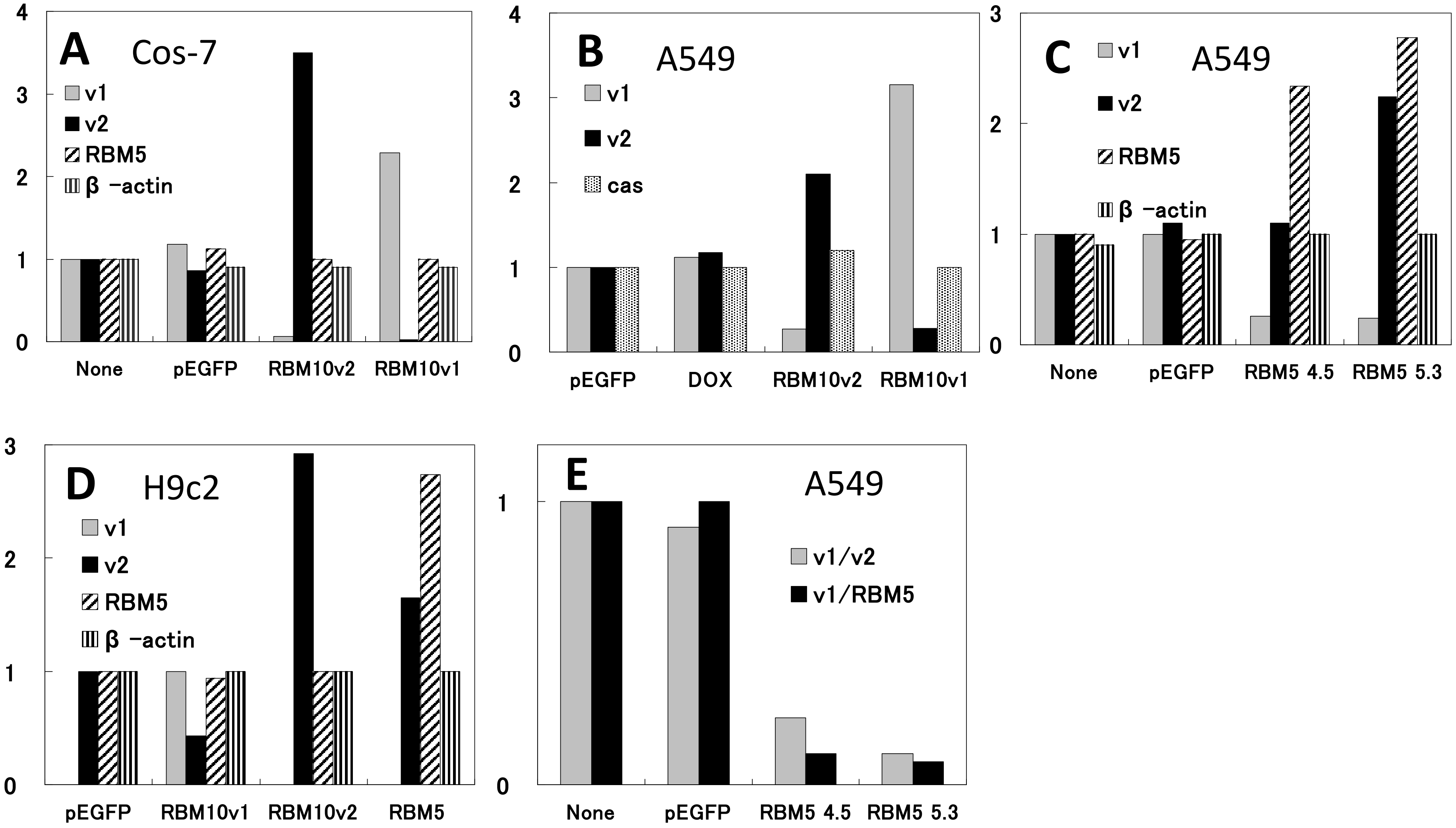
