## supplement S2 for "RBM10 variants and RBM5 involve in alternative splicing of RBM10v1 pre-mRNA: RBM10v1 includes its own exon 4, but RBM10v2 and RBM5 skip it"

```
                          1        10        20         30        40       49
human RBM10v1 RRM1   (1) IVMLRMLPQAATEDD I RGQLQSH - GVQAREVRL MRNKS SGQSRGFAFVEF
human RBM10v2 RRM1   (1) ------------------------------E I RGQLQSH - GVQAREVRL MRNKS SGQSRGFAFVEF
   human RBM5 RRM1   (1) T IMLRGLP I T ITES D I REMMESFEGPQPADVR LMKRKT - GVSRGFAFVEF
   human RBM6 RRM1   (1) – IRLS GVPEDAT KEE I L NA FRTPDGMPVKNLQL KEYNT - GYDYGYVCVEF
         Consensus   (1)   IMLRGLP    A TEDD I RGQLQSHDGVQAREVR LMRNKSSGQS RGFAFVEF
```

```
                          50         60       70      78
human RBM10v1 RRM1  (50) SHLQDATRWM EANQHSLNI LGQKVSMHYS   % similarity to RBM10v1 RRM1
Human RBM10v2 RRM1  (36) SHLQDATRWMEANQHSLNI LGQKVSMHYS   98.4% (identity)
   human RBM5 RRM1  (50) YH LQDATSWM EANQKKLVI QGKHIAMHYS  59.5% (identity), 68.4% (positive identity)
   human RBM6 RRM1  (49) S L LEDAI GCM EANQGTLMIQDKEVT LEY -  29.5% (identity), 44.9% (positive identity)
         Consensus  (51) SHLQDATRWMEANQHSL NI QGQKVSMHYS
```

```
                          1        10        20        30         40       50
human RBM10v1 RRM2   (1) T I I LRNLNPHSTMDSI LGALAPYAVLSSSNVRV I KDKQTQ LNRGFAF IQL
human RBM10v2 RRM2   (1) TI I LRNLNPHSTMDSI LGALAPYAVLSSSNVRV I KDKQTQ LNRGFAF IQL
   human RBM5 RRM2   (1) TI I LRNI APHTVVDSI MTALSPYASLAVNN I RL I KDKQTQQNRGFAFVQL
   human RBM6 RRM2   (1) - IMLKRI YRSTPPEVI VEVLEPYVRLT TANVR I I KNRTGPMGHTYGF IDL
         Consensus   (1) T I I LRNINPHSTMDSI LGALAPYAVLSSS NVRVI KDKQTQL NRGFAFI QL
```

```
                          51       60        70         82
human RBM10v2 RRM2  (51) S T I VEAAQLLQI LQALHPPLTIDGKTI NVEFA    %similarity to RBM10v1 RRM2
human RBM10v2 RRM2  (51) S -T I E AAQLLQI LQALHPPLTIDGKTI NVEFA   98.8% (identity)
   human RBM5 RRM2  (51) SSAMDASQLLQ I LQSLHPPLKIDGKTI GVDFA    69.5% (identity), 87.8% (positive identity)
   human RBM6 RRM2  (50) DSH AEALRwK I LQNLDPPFSIDGKMV--------  32.9% (identity), 56.1% (positive identity)
         Consensus  (51) S S I - EAAQLLQ I LQALHPPLTIDGKTI NVEFA
```
